# Synchrony is a robust iEEG biomarker for antiseizure medication load in epileptic patients

**DOI:** 10.64898/2026.01.09.698669

**Authors:** Devin Ma, Nina J. Ghosn, Carlos A. Aguila, Quy Cao, Erin C. Conrad, Brian Litt

## Abstract

With new, minimally invasive continuous EEG and wearable monitoring systems for epilepsy nearing regulatory approval, there is a rush to determine how to make these devices most useful to patients. One application is to track biomarkers of antiseizure medication (ASM) levels to help manage seizure risk and side effects. In this paper, we validate one such biomarker, phase synchronization, during medication taper and presurgical evaluation in a cohort of 80 consecutive patients recorded with intracranial EEG (iEEG) at the University of Pennsylvania. While previous investigators have demonstrated that synchrony is negatively correlated with ASM load at the resolution of 1 day, we hypothesize that synchrony continuously tracks and can predict ASM load on a clinically useful time scale. We test this hypothesis using a pharmacokinetic model generating continuous ASM load values previously published by our group and correlate it with continuous synchrony values derived from the iEEG. We use a linear mixed effect model to examine the relationship between ASM load, synchrony, vigilance, and other time dependent factors. We use dominance analysis to rank the relative importance of predictors based on their influence on ASM load. We find that synchrony not only can predict ASM load but is also the most significant predictor. Our study highlights the potential of synchrony as a biomarker for ASM load, and its utility in an ambulatory implantable device that can alert patients to conditions lowering their medications (e.g., adherence, drug interactions, generic change, pregnancy, etc.) or potential medication toxicity. We propose that measures like synchrony, packaged in an appropriate interface, could bring substantial value to ambulatory epilepsy management devices.

## Introduction

For most individuals with epilepsy, antiseizure medications (ASMs) are the first line of treatment. Despite this, only two-thirds of patients may achieve seizure freedom with ASM alone, underscoring the need for better management strategies for the remaining drug-resistant population. Effective management of epilepsy, particularly in drug-resistant cases, often requires precise modulation of ASM load, as missing even a single dose can trigger breakthrough seizures that significantly impact quality of life.^1^

Antiseizure medications function in part by stabilizing neuronal excitability and reducing hypersynchronous neuronal discharges, which are hallmarks of epileptic seizures. In epilepsy monitoring units (EMUs), clinicians commonly taper ASMs in drug-resistant patients to provoke seizures for surgical planning purposes. This controlled manipulation of ASM load offers a unique opportunity to study the effects of ASMs on brain network dynamics, which remain underexplored.

There is growing evidence that suggests epilepsy is a disease of brain networks,^2,3^ whereby seizures are characterized as aberrant hypersynchronous discharges and abnormal excitatory/inhibitory (E/I) balance.^4^ ASMs work in part by managing the E/I balance and thereby suppressing the occurrence of hypersynchronous activity.^5^ However, little has been done to quantify changes in network dynamics in response to varying ASM load.^6,7^ Establishing the link between ASMs and brain network changes can enhance our understanding of the mechanisms of both the disease and medications, while also serving as the basis for a biomarker of seizure risk due to changing ASM loads.

Increasingly, implantable neurostimulation systems, such as the Responsive Neurostimulation (RNS) device (NeuroPace, Mountain View, CA), hold promise for stimulating the seizure onset zone (SOZ) to prevent seizures.^8^ However, these devices rarely consider the effect of antiseizure medication load on EEG signals, even though a patient’s propensity to seize likely depends, in part, on whether the patient is adherent to their medication regimen. It is common for patients who miss a dose of medications to experience more debilitating seizures.^9^ Other conditions that affect ASM levels include decreased availability of medications after a generic switch, drug interactions, pregnancy, or potential degradation of medications that might either decrease or increase levels to the point of causing side effects. A reliable EEG biomarker for ASM level could be an invaluable aid to disease management and bring important value to EEG monitoring devices beyond counting seizures. Of course, many of these devices do not utilize intracranial EEG. We have chosen to do this first study using iEEG because of its superior signal quality and decreased artifact, before moving to noisier, less invasive signals.

Brain-wide synchrony has emerged as a promising network metric to correlate with medication load. Previous studies suggest that phase synchronization calculated from iEEG is negatively correlated with medication load.^10–12^ Over a patient’s stay in the epilepsy monitoring unit with iEEG monitoring and ASM tapering, days with the highest overall ASM load were found to have the lowest iEEG synchrony. However, these studies use daily averages of medication load, while medication load can vary greatly during taper, according to each medication’s pharmacokinetic profile. As such, analysis on a finer temporal granularity is needed to establish whether synchrony can be employed as a potential biomarker of ASM load on a time scale that has greater clinical utility.

Here, we hypothesize that synchrony not only tracks ASM load but can also serve as an iEEG biomarker for medication load. Below, in a series of analyses, we correlate real-time estimates of ASM load to iEEG synchrony while controlling for potential confounding factors.

## Materials and methods

### Patient cohort

We retrospectively studied 80 consecutive patients with drug-resistant epilepsy who underwent iEEG monitoring in the EMU for presurgical planning at the Hospital of the University of Pennsylvania (HUP) between January 2015 and October 2021. These patients have well-documented and previously validated seizure times, seizure categorizations, and exact times and doses of ASM administrations. We obtained informed written consent from each patient. Data collection was approved by the Institutional Review Board at the Hospital of the University of Pennsylvania. More details on patient demographics can be found in Table S1 in Supplementary Material.

### iEEG recording and processing

Patients undergoing iEEG monitoring were implanted with linear cortical strips, cortical grid arrays, and linear depth electrodes. Sampling rates of recordings ranged from 256 Hz to 1024 Hz. We followed standard pre-processing techniques for iEEG data, illustrated in Fig. 1A. We first divided each patient’s full recording into 2-minute segments for tractable computation. Next, we performed automated artifact rejection on each electrode of each 2-minute segment and discarded electrodes with excessive high voltage deviations, high variance, and excessive line noise, following previously validated processes.^13,14^ We then re-referenced each electrode to a common average reference (CAR). Finally, we used a notch filter (order 2 Butterworth filter, band stop with low-pass 59 Hz and high-pass 61 Hz) to remove 60 Hz line noise.

**Figure 1.**
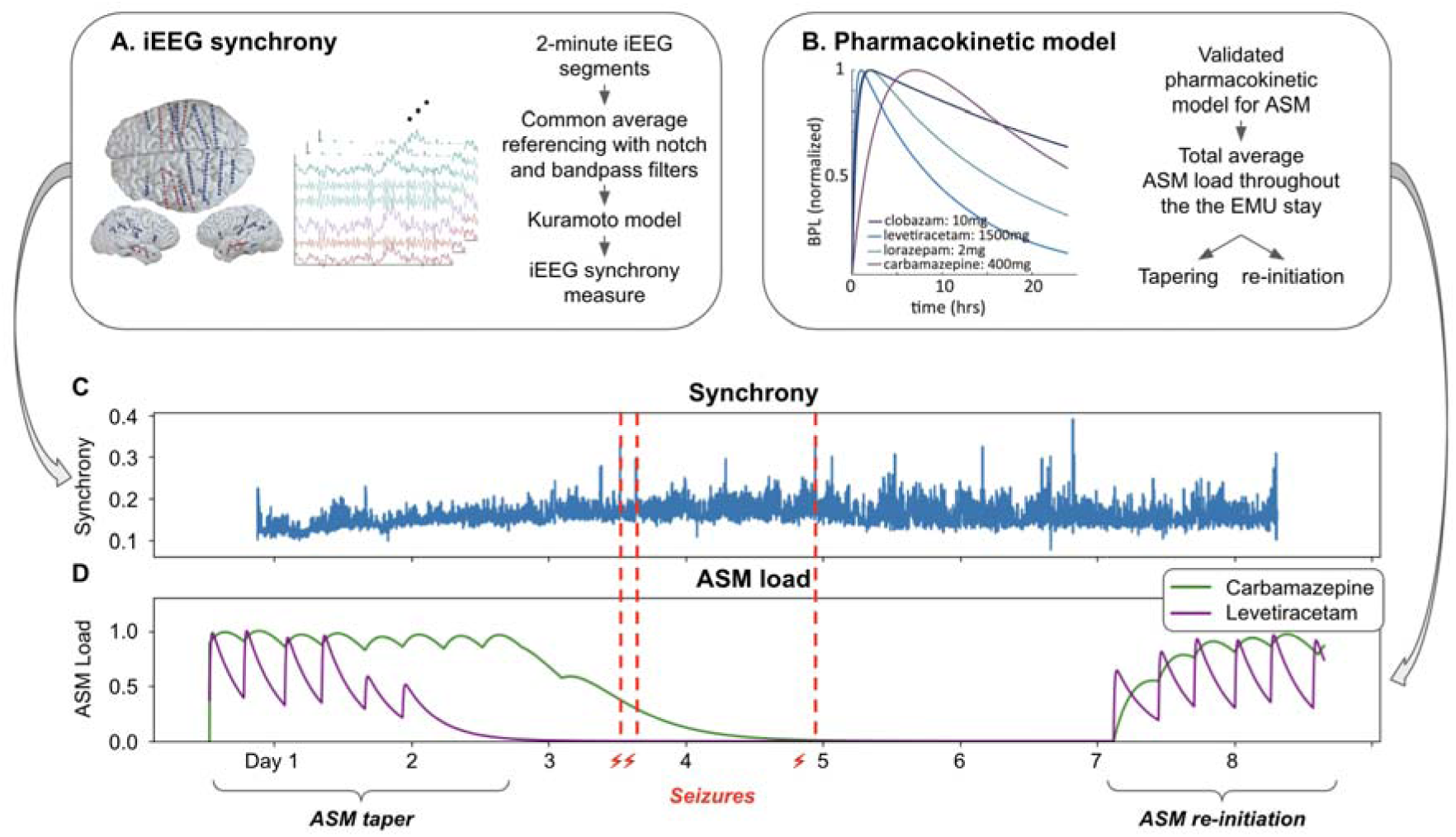
Toolkit and analysis pipeline. **(A)** For each patient, we compute intracranial EEG synchrony measures across all electrodes using the Kuramoto model. **(B)** We use a previously validated pharmacokinetic model for antiseizure medication (ASM) in the epilepsy monitoring unit (EMU) setting to model the continuous load of each ASM during the patients’ EMU stay. We also determine when tapering and re-initiation of medications occurs. We juxtapose an example patient’s (HUP 142) iEEG synchrony time series in **(C)** with the same patient’s ASM load curves in **(D)**, where we annotate the ASM tapering, the subsequent seizures, and re-initiation of the medications towards the end of the EMU stay.

### iEEG synchrony calculation

Synchrony measures the extent of phase locking between two or more oscillatory signals. When computed as phase synchronization using the Kuramoto model, synchrony has been used in previous literature as a network metric to characterize the global synchronization of neural oscillations and global levels of cortical excitability.^5,11,12,15–17^ To calculate synchrony, we first obtain the angle (arctan) of the Hilbert transform *H*(*x*) of the pre-processed iEEG data *F_i_*(*t*), resulting in complex-valued analytical signals. The real part represents the original signal, while the imaginary part represents the phase information. Then, *r_t_*, the average phase synchronization in each time segment *t* is calculated by taking the mean of the complex exponentials of the phase angles across electrode channels at each time point. The average synchrony across all electrodes in each time segment, *R*, was then calculated. This was done for 2-minute time segments *t* across the EMU stay. The average synchrony of each segment is then the average of all Kuramoto order parameters. This calculation is summarized by the set of equations below:

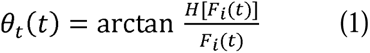

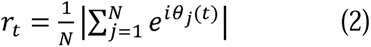

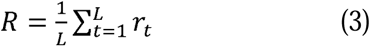

### Alpha-delta ratio as wakefulness measure

Previous studies have shown that circadian patterns, particularly wakefulness and sleep, can affect iEEG metrics.^15,18–23^ To account for the effect of wakefulness on synchrony, we computed the ratio of power in the alpha (8-13 Hz) and delta (1-4 Hz) frequency bands. This alpha-delta ratio was computed for each time segment and normalized across all segments for a patient. Normalized alpha-delta ratio has been previously established as a measurement of wakefulness^24–26^ and has been validated in this patient cohort for another study within our group to represent wakefulness state.^14^

### ASM administration in the EMU

Administration time and dosing information of ASMs, including both scheduled medications and rescue doses of benzodiazepines, were extracted from the patients’ electronic health records. The patient’s medication regimen during the first day of EMU admission was used to determine the initial dosage levels.

### Pharmacokinetic model of ASM load

ASMs are absorbed and decay in the bloodstream in a predictable manner and were modelled using a previously validated first-order pharmacokinetic model to generate continuous curves of blood plasma levels for each medication.^27^ The model considers the bioavailability of each drug, volume of distribution, and absorption and elimination half-lives. The overall load of ASM was calculated as the average of the normalized individual curves generated by the model. To ensure that the ASM load accurately reflects the steady-state pharmacodynamics of the regularly administered medications, we excluded rescue medications like lorazepam in the total load average. Each curve was normalized to the maximum ASM load during the EMU stay. Average ASM load estimates were correlated with iEEG synchrony values in subsequent analyses. Using this model of ASM load, we subsequently define tapering and re-initiation of ASMs in the EMU and find the exact time of these events to test their effect on iEEG synchrony levels (Fig. 1B).

### ASM taper

In later sections, we will leverage the full cohort’s consecutive iEEG recording and ASM curves to investigate the continuous relationship between ASM load and synchrony in a linear mixed model. First, we pursued a simpler case—in the patients who had *significant* ASM taper, how did synchrony change in response to the significant reduction in total ASM load. This is because while all 80 patients had fluctuations in ASM load, only some patients experienced significant ASM taper (Fig. 1C-D), typically during the first two days of the EMU stay. This allows us to test the relationship between ASM load and synchrony in a pre-ictal environment, isolating the potential effect of seizures on synchrony.

Here, we defined such ASM taper event as when the total ASM load drops below 50% of the maximum total load since the start of the EMU stay. We saw this significant medication taper in 16 patients (majority with left-sided SOZ laterality and temporal SOZ localization, see Table S1 in Supplementary Material).

### ASM re-initiation

After seizures are induced with ASM taper, clinicians usually re-initiate admission ASMs for patients towards the end of the EMU stay prior to discharge or initiate a new therapy (see example in Fig. 1C-D). This gives us the opportunity to study whether synchrony responds to ASM re-initiation, as well as taper. As in ASM taper, we again sought to first find patients with significant ASM re-initiation at the end of the EMU stay to test how synchrony responds to such events, even though there may be other instances of total ASM levels increasing, which will be incorporated into the linear mixed model later.

Here, we defined such re-initiation event as when the total ASM load increases by at least 50% from the minimum load after their last recorded seizure in the EMU stay. Some patients in our cohort did not have ASM re-initiated, while others had limited data iEEG data after their last seizure. We found that nine patients (majority with left-sided SOZ laterality and temporal SOZ localization, see Table S1 in Supplementary Material) had ASM re-initiation in a way that fit the above criteria, all within the final 48 hours of their EMU stay.

### Effect of ASM taper and re-initiation on synchrony

To study how iEEG synchrony responds to changes in ASM load, we first tested how synchrony changes after ASM taper. To do this, we compared the iEEG synchrony values 12 hours before and 12 hours after the taper. To isolate the effects of seizure activity, we only included patients who were seizure-free within the 12 hours after taper. We used Mann-Whitney U tests with Bonferroni correction to compare the synchrony values before and after tapering. We then repeated this analysis for the nine patients with ASM re-initiation to test if and how synchrony responds to re-initiation.

### Isolating the effect of seizures on synchrony

To establish the relationship between synchrony and ASM load, it is important to rule out the possibility that the relationship between synchrony and ASM load is driven by seizures (i.e., how close the time we measure synchrony was to a seizure).

To do this, we examined data from a 90-minute pre-ictal window, specifically from 120 to 30 minutes before each seizure. We first identified valid seizure events, ensuring that no seizures occurred within 120 minutes of each other, as closely spaced seizures could distort synchrony measurements. For each valid seizure, we calculated the average synchrony and ASM load in the defined pre-seizure window. This approach allows us to assess how synchrony responds to varying levels of ASM load, independent of the immediate effects of a seizure. To quantify the relationship between ASM load and synchrony, we then performed ordinary least squares (OLS) regression analyses using pre-ictal synchrony values as the dependent variable and ASM load as the independent variable.

### Continuous relationship between ASM load and synchrony

Having tested how synchrony responds to ASM taper and re-initiation, we next investigate the continuous relationship between synchrony and ASM load throughout the EMU stay. We hypothesize that synchrony continuously tracks ASM load beyond the periods immediately before and after ASM tapering and re-initiation.

We identified 68 patients (see Table S1 in Supplementary Material) who had fewer than 10 seizures with the time between successive seizures being at least 2 hours to isolate the effect of seizure dynamics on synchrony. We then applied a linear mixed model using restricted maximum likelihood (REML) estimation to the 68 patients’ iEEG recordings of entire EMU stay to examine the relationship between total ASM load (as the dependent variable), and other covariates (synchrony and alpha-delta ratio), while accounting for both fixed and random effects. Since some patients experienced multiple seizures, we included time dependent covariates (time since previous seizure, time since EMU admission) in the model.

Next, to evaluate the relative importance of different predictors in our mixed-effects model, we performed a bootstrap resampling analysis with 1,000 iterations to quantify the variability in the coefficients of four key predictors: synchrony, time since previous seizure, EMU monitoring time, and alpha-delta ratio.^28^ We generated 1,000 bootstrap samples by sampling with replacement from the full dataset. Each bootstrap sample had the same size as the original data, and it was used to fit a new mixed-effects model. For each bootstrap sample, we fitted the same linear mixed-effects model as the one in the previous section to predict the ASM load with patient-specific random intercepts included to account for within-subject variability. After fitting each model, we extracted the estimated coefficients for the four predictors and stored them for further analysis. This procedure was repeated for each bootstrap sample, resulting in a distribution of coefficients for each predictor across all 1,000 bootstrap iterations. We calculated the standard deviation of the coefficients for each predictor across the bootstrap iterations as a measure of their variability and relative importance. Predictors with higher standard deviations were considered more variable and potentially more impactful in the model.

To ensure that coefficient estimates were comparable across predictors and that variability in bootstrap estimates reflected meaningful differences rather than differences in scale, we standardized all continuous variables prior to model fitting.

### Synchrony’s robustness to electrode sampling

We assessed our synchrony’s robustness to electrode sampling by calculating synchrony for each patient using varying electrode percentages, from 10% of total number of iEEG electrodes available to 100% in 10% increments. Electrodes were randomly selected for each subsample. Synchrony was computed for 50 evenly spaced 2-minute intervals pre-seizure. We then calculated Pearson correlations between subsampled and full-sample synchrony values. Median values and interquartile ranges were reported across all patients.

## Results

### Synchrony increases as ASM is tapered

To understand the relationship between synchrony and ASM load, we first tested how synchrony changes after ASM tapering in the EMU. We found that all 16 patients that had ASM tapered had a statistically significant increase in the synchrony value after tapering (*P <* 0.0001, Bonferroni-corrected, see Fig. 3A). This indicates that synchrony values increase significantly as ASMs are tapered.

We further hypothesize that the increase in synchrony values after taper happens gradually as the patient’s medication load decreases. To test this, we computed the average daily synchrony value starting from the first day of the patient’s EMU stay until their first seizure. We observed that in some patients (e.g., HUP 142), there is a steady daily increase in average synchrony values in the days after EMU admission leading up to the first seizure induced by ASM tapering (Fig. 2A). The statistically significant distribution shift in synchrony values before and after tapering for this patient is shown in Fig. 2B. This suggests that synchrony may be a useful proxy to ASM load and can be sensitive to ASM tapering.

**Figure 2.**
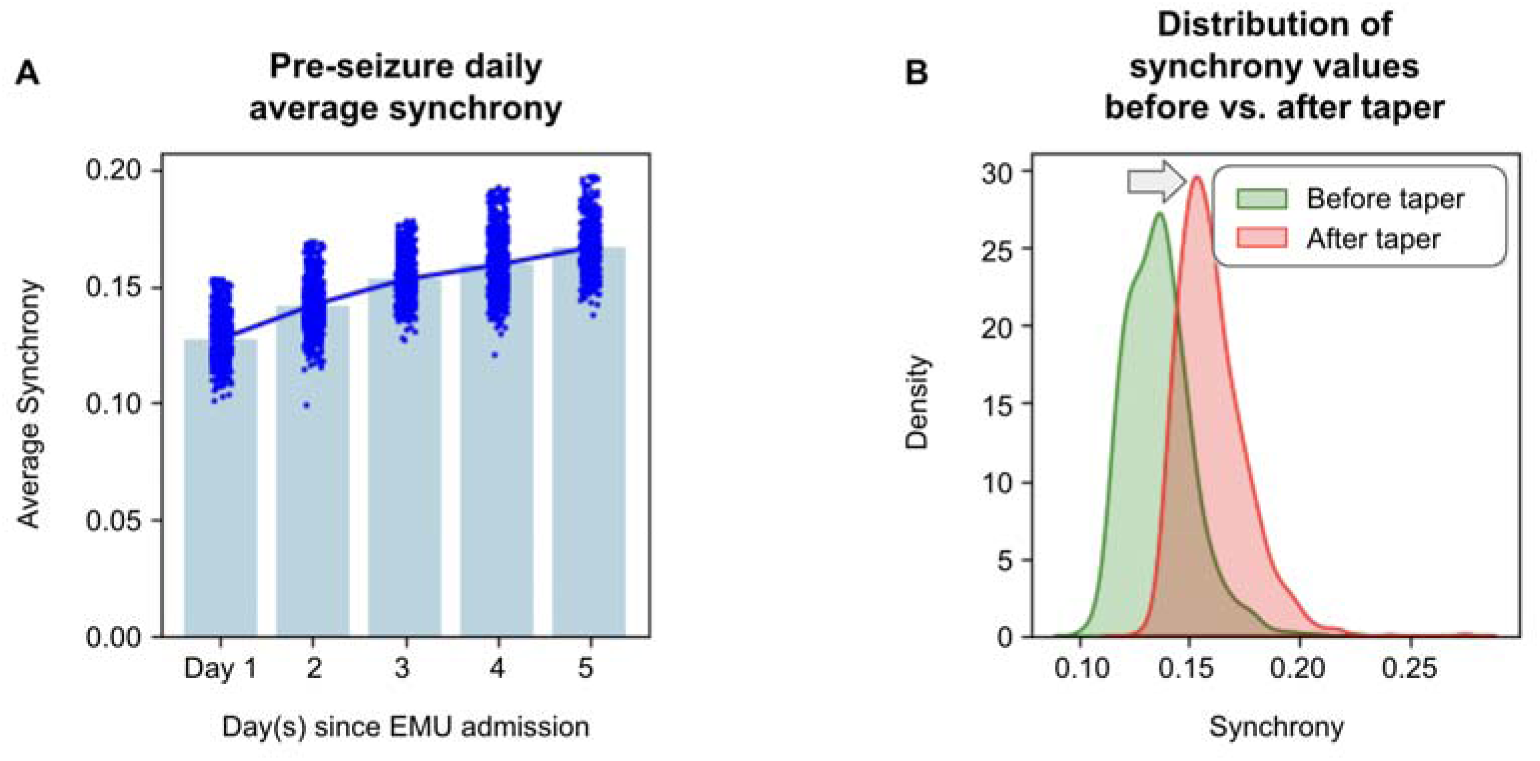
Synchrony increases as ASM is tapered in a single patient. **(A)** Daily average synchrony increases as ASM is tapered on each day before a seizure in the EMU in an example patient. Each blue dot represents the synchrony value of a 2-minute iEEG segment. **(B)** Distribution of synchrony values significantly shifts right after ASMs are tapered.

### Synchrony may decrease as ASM is re-initiated

If synchrony increases as ASMs are tapered, we hypothesize that the inverse holds too, that synchrony decreases as ASMs are re-initiated. Out of the nine patients that had ASM re-initiated towards the end of the EMU stay, we found that five patients had a statistically significant decrease in the synchrony value after ASM re-initiation at the *P <* 0.001 level (Bonferroni-corrected), two patients showed significance at the *P <* 0.01 level (Bonferroni-corrected), and 2 others were not statistically significant (*P >* 0.05), as shown in Fig. 3B. The Bonferroni correction was applied to account for multiple comparisons, with an adjusted significance threshold of *P <* 0.0056 for *P <* 0.05, *P <* 0.0011 for *P <* 0.01, and *P <* 0.00011 for *P <* 0.001, calculated based on 9 comparisons. Taken together, this indicates that, at least in some patients, synchrony is responsive to not only tapering but also re-initiation, suggesting that it could be a useful biomarker for changes in ASM load.

**Figure 3.**
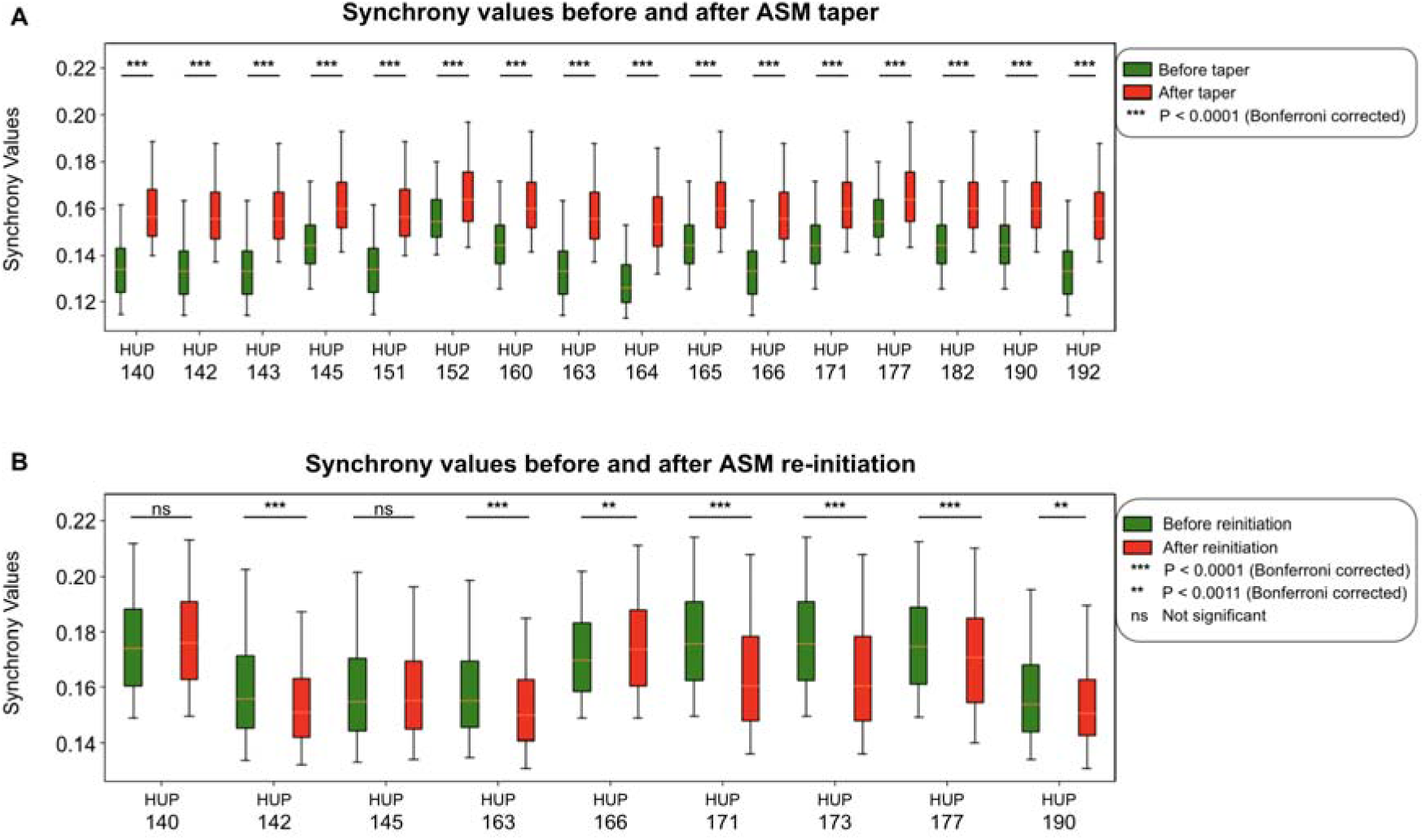
Synchrony increases after tapering and decreases after ASM re-initiation. **(A)** 16 patients with ASM tapered at the start of the EMU stay had statistically significant increase in synchrony values after tapering. **(B)** seven out of the nine patients who had ASM re-initiated towards the end of EMU stay had statistically significant decrease in synchrony values.

### Pre-ictal synchrony tracks ASM load instead of seizure activity

One potential confounder in the relationship between ASM load and synchrony in this setting is seizures, which may drive increases in synchrony. Hence, we next asked if synchrony during the period right before seizures is correlated with ASM load. If we see a relationship between pre-ictal synchrony and ASM load, and if seizures that occur at similar ASM loads have similar pre-ictal synchrony values within a patient, then this would support the hypothesis that synchrony can be a biomarker for ASM load.

We performed an OLS regression using pre-ictal synchrony values as the dependent variable and ASM load as the independent variable (Fig. 4A). The OLS regression yielded an *R*^2^ value of 0.030, indicating that 3% of the variability in synchrony can be explained by ASM load in the baseline period. The coefficient for ASM load was −0.0184 (*P* = 0.032), demonstrating a statistically significant negative relationship between synchrony and ASM load. The constant term in the model was 0.1253 (*P* < 0.001), representing the average synchrony when ASM load is zero. The model’s adjusted *R*^2^ value of 0.024 reflects that this relationship holds, though the effect size is small. Importantly, the negative coefficient for ASM load suggests that even during stable, non-seizure periods, lower ASM levels are associated with higher synchrony. This finding supports our hypothesis that ASM load suppresses synchrony, consistent with its role in reducing neural excitability. Despite the modest explanatory power of the model, the significant *P*-value for ASM load highlights the robustness of synchrony as a potential biomarker for ASM load. This effect persists even after accounting for patient-specific variability, underscoring synchrony’s potential utility in predicting changes in medication levels.

**Figure 4.**
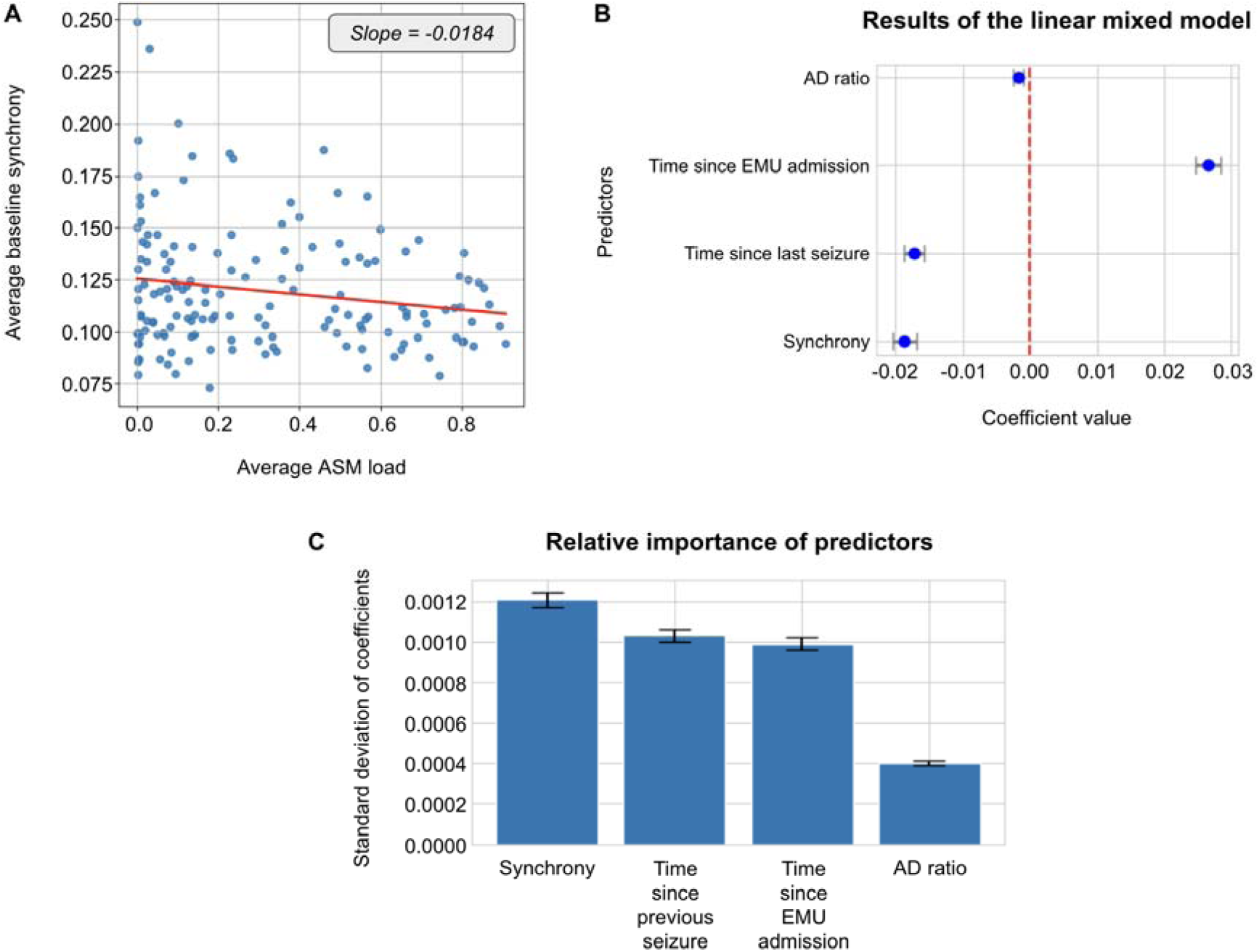
Synchrony tracks ASM load continuously throughout the EMU stay. **(A)** Baseline pre-ictal synchrony and ASM load from all seizures in all patients are negatively correlated. **(B)** Coefficients from the linear mixed model predicting ASM load showing synchrony has a significant negative correlation to ASM load. **(C)** Synchrony captures more variance in ASM load than any other variable we considered, with the highest coefficient variability (mean ± std, 0.001175 ± 0.000037).

### Synchrony tracks ASM load continuously in a linear mixed model

Next, we tested the continuous relationship between synchrony and ASM load throughout the EMU stay. Our hypothesis is that synchrony continuously tracks ASM load beyond the periods immediately before and after ASM tapering and re-initiation. We applied a linear mixed model using restricted maximum likelihood (REML) estimation to examine the relationship between the total ASM load as the dependent variable and other covariates (synchrony, time since previous seizure, time since EMU admission, and alpha-delta ratio), while accounting for both fixed and random effects. As shown in Fig. 4B, the mixed-effects model shows a significant inverse relationship between synchrony and antiseizure medication load (Coef. = −0.019, *P* < 0.001). This indicates that as synchrony increases, ASM load decreases. Specifically, synchrony had a highly significant negative coefficient, confirming that medication load has a strong, consistent effect on synchrony across patients. Additionally, the effect of time since previous seizure (Coef. = −0.017, *P* < 0.001), time since EMU admission (Coef. = 0.027, *P* < 0.001), and alpha-delta ratio (Coef. = −0.002, *P* < 0.001) were also significant. While these variables remain statistically significant, their coefficients suggest comparatively smaller—but still measurable—effects on ASM load.

### Synchrony accounts for the most variance of ASM load

Following the linear mixed model analysis, we asked which of the covariates explained the most variance in ASM load. To assess this, we performed a bootstrap resampling analysis with 1,000 iterations, evaluating the variability and relative importance of four key predictors—synchrony, time since previous seizure, time since EMU admission, and alpha-delta ratio—in a mixed-effects model. Higher standard deviations of the coefficients indicate greater variability and potential impact on ASM load. As shown in Fig. 4C, synchrony exhibited the highest coefficient variability (0.00118 ± 0.00004), indicating it is the most influential predictor in the model. In contrast, time since previous seizure (0.00096 ± 0.00003), time since EMU admission (0.00095 ± 0.00003), and alpha-delta ratio (0.00039 ± 0.00001) showed considerably lower variability, suggesting that their contributions to ASM load were comparatively minor.

These results indicate that broadband synchrony has a more substantial impact on predicting ASM load than the other factors considered. The small standard errors for the latter three predictors reflect the high precision of their coefficient estimates, though their relative influence on ASM load was minimal.

### Synchrony could be a robust biomarker for ASM load

Intracranial EEG monitoring in the EMU setting often involves over 100 electrodes implanted in the seizure onset zone. For example, in our patient cohort, patients had on average 120 electrodes. While this enables better spatial sampling of cortical dynamics to facilitate seizure diagnosis and localization, the practicality of storing and processing large amounts of continuous iEEG data over a long period of time becomes limited outside of the EMU setting. We are interested in assessing the utility of synchrony as a biomarker on a smaller set of electrodes.

We first notice that using fewer electrodes typically results in higher synchrony values (Fig. 5A) due to how synchrony is computed (it’s much more likely to have higher phase locking value when there are fewer oscillators). We predicted that there was a minimum percentage of subsampled electrodes that would be sufficiently similar to synchrony sampled from all electrodes. Across our cohort, the median Pearson correlation increases as the percentage of electrodes are sampled (Fig. 5B). The largest increase in correlation occurs from 10% to 20% of sampled electrodes. At 40% sampling, the median Pearson correlation (median [IQR]: *r* = 0.85 [0.63-0.90]) is strongly correlated to full sampling. These results imply that a subsampled network achieves similar synchronization values and patterns across time in this setting, and possible in outpatients with implantable devices.

**Figure 5.**
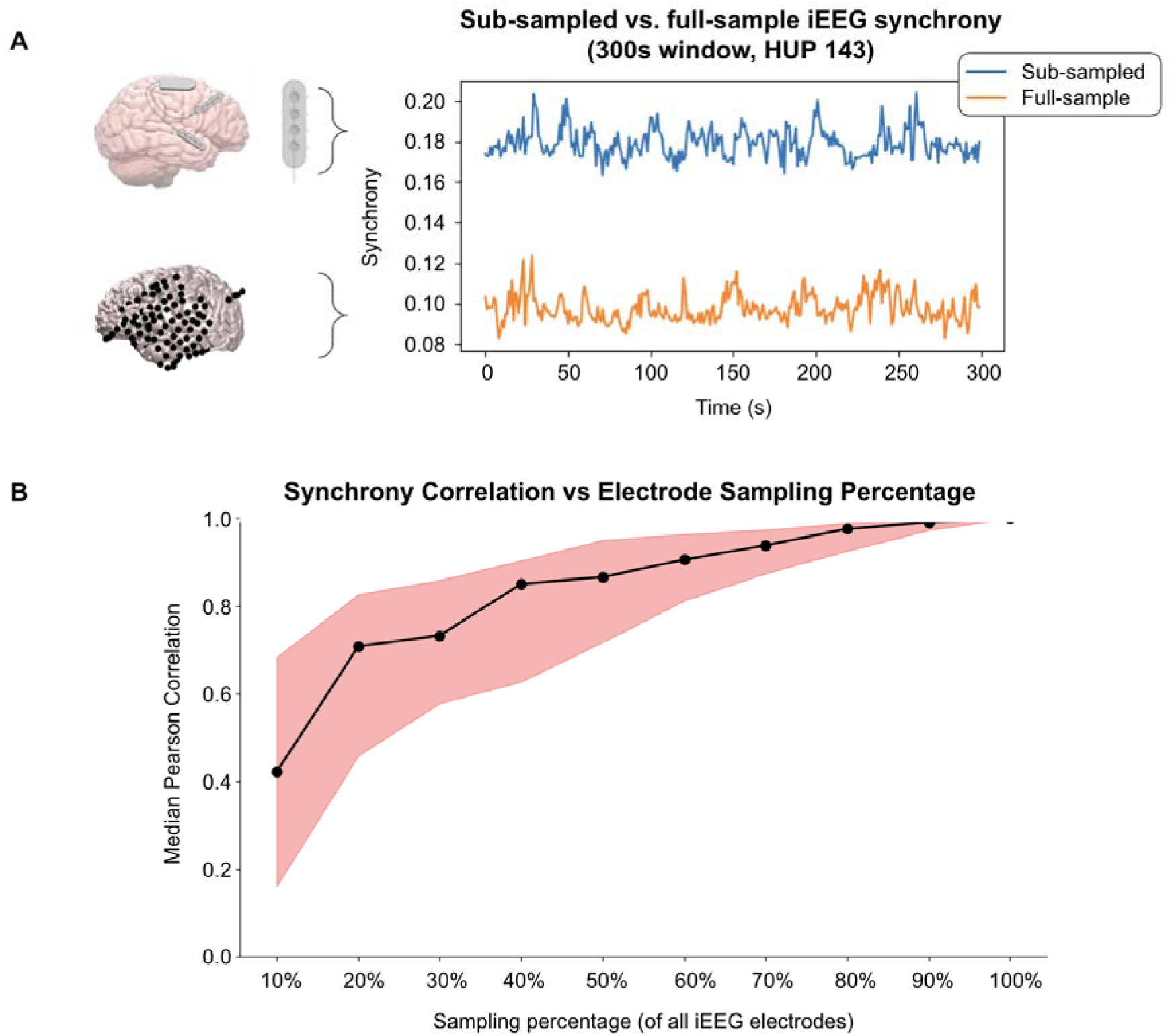
Synchrony can be a robust biomarker for ASM load. **(A)** Schematic showing a 300-second window of sub-sampled synchrony and full-sample synchrony of one example patient. **(B)** The median Pearson correlation increases as the percentage of electrodes sampled increases. The largest increase in correlation occurs from 10% to 20% of sampled electrodes. At 40% sampling, the median Pearson correlation (median [IQR]: *r =* 0.85 [0.63-0.90]) is strongly correlated to full sampling.

## Discussion

In this study, we observed that synchrony can be a robust iEEG biomarker for ASM load. We identified patients who had ASMs tapered at the beginning of their EMU stay and showed that synchrony increased after ASM load was reduced. We then identified patients who had ASMs re-initiated at the end of their EMU stay and showed that in some patients, synchrony decreased after ASM load was increased. This led us to believe that synchrony may track ASM load in general and can be a useful measure of cortical dynamics under the influence of ASMs. To study the relationship, we employed a linear mixed model and observed that synchrony has a significant negative correlation with ASM load. Since wakefulness has been shown to influence various iEEG measures,^14^ we included alpha-delta ratio as a covariate in the mixed model to represent wakefulness. The results of the linear mixed model showed that not only is synchrony negatively correlated with ASM load, but it is the most important predictor and captures more variance of ASM load than any other predictor we considered.

### Implication for seizure dynamics

The findings of this study align closely with the theoretical model of self-organized criticality, which posits that the brain operates near a critical point, characterized by balanced excitatory and inhibitory neural activities.^29,30^ Operating near this critical point is thought to facilitate optimal responsiveness to external inputs while avoiding runaway excitatory events like seizures.^17,31–34^ Epileptic seizures are often considered “supercritical” events in which the brain transitions from a subcritical or near-critical state to a state of excessive synchrony and hypersynchronous discharges.^35^ Seizures represent a breakdown of the brain’s normal inhibitory controls, where neural networks fire in an overly synchronized manner, disrupting normal function.^16,36^ In this context, ASMs are thought to modulate neural networks by maintaining a subcritical state, thereby preventing the brain from tipping into pathological synchrony and seizure activity.^15^

In this study, we demonstrated that iEEG synchrony serves as a robust biomarker for ASM load. As ASM load is reduced, iEEG synchrony increases, indicating a shift toward higher cortical excitability. This gradual increase in synchrony may signal a drift toward a supercritical state, eventually culminating in seizure onset. This supports the hypothesis that epileptic seizures arise after a period of heightened seizure risk, marked by runaway excitatory and hypersynchronous activity in the cortical networks.^37^

Further evidence supporting the idea of brain criticality comes from the inverse relationship observed between ASM re-initiation and iEEG synchrony. As ASMs are reintroduced, synchrony decreases in a manner suggesting that ASM helps restore the brain’s subcritical, balanced state, reducing cortical excitability and preventing seizures. This observation resonates with findings from studies that showed that ASMs push the brain into a subcritical dynamic state, reducing the risk of supercritical, seizure-like events. The criticality framework also explains why synchrony can serve as an early indicator of seizure risk.^38^ As the brain nears criticality, small fluctuations in network dynamics, like changes in synchrony, may precede larger transitions, such as seizures. This opens the door for the use of synchrony as a biomarker not only for ASM load but also for seizure risk, which we can subsequently control by manipulating ASMs, though these two variables are linked.

### Utility in implantable recording devices

We were also interested in assessing the utility of synchrony as a biomarker outside of the EMU setting. In our dataset, iEEG monitoring often involves over 100 electrodes per patient, while the latest implantable devices typically have under 10 electrodes. Our results show that while the magnitude of synchrony depends on the number of electrodes (e.g., with fewer electrodes it’s much easier to have larger synchrony), there is meaningful correlation between the synchrony computed with all >100 electrodes and a subsampled calculation. Furthermore, in some patients, as the number of electrodes used in computing synchrony increases, the correlation with full sample synchrony stops increasing (i.e., the curve plateaus). Since electrodes further removed from the parenchyma, such as subdural, subgaleal or scalp electrodes, record from larger regions of tissue, it may be that fewer of these electrodes may yield even better measures for clinical use, than intracranial electrodes that record from much more confined regions. Our findings suggest that it may be possible to utilize synchrony as a biomarker in a device that has under 10 electrodes, although future work is needed to validate it in the outpatient setting, and with different types of sensors in minimally invasive devices.

### Enabling new capabilities in implantable devices

Epileptic seizures are often unpredictable both in timing and severity. For outpatients, there are many factors that can aggravate seizure risk. These factors can be environmental factors such as excessive heat, or factors that patients can control, such as alcohol consumption or missing medications. Our study demonstrates how manipulating one of the most prominent factors, ASM load, can alter cortical dynamics and ultimately affect seizure risk.

If validated, new iEEG biomarkers such as synchrony could create a new mode of “patient-in-the-loop” paradigms in an outpatient setting with wearable and potential closed loop devices, such as the NeuroPace RNS and other devices currently in development. Currently, responsive neurostimulation devices work by detecting changes in the iEEG signals and stimulating the epileptogenic regions accordingly. This is a “closed loop” model since an algorithm determines the timing and intensity of stimulation without the patient’s interference. While this is undoubtedly an advance from earlier generations of neurostimulation, this current paradigm removes the consideration of factors the patient can actively manage to reduce seizure risk, and the need for particular instances of stimulation. If the device can detect changes in iEEG biomarkers such as synchrony, it can then instruct patients to take their prescribed dose of medications missed, proactively manage seizure risk.

### Limitations and future directions

Our study does not present without limitations. First, the placement and number of intracranial EEG electrodes varied between patients, as they were implanted based on clinical necessity to localize epileptogenic zones. This variability in electrode coverage means that we may have missed certain cortical dynamics that were not captured, potentially affecting our findings related to synchrony and ASM load modulation.^39^ However, most implants are placed in the hypothesized seizure onset zone, a strategy that is also followed with implantable devices.

Second, while we identified synchrony as a robust biomarker for ASM load, the presence of circadian influences on iEEG biomarkers may not have been fully controlled for. Previous research indicates that inter-ictal features and seizure activity follow circadian rhythms.^14,22,40^ Though we attempted to account for these variations through the inclusion of a validated measure of sleep, other circadian modulation could introduce additional confounding effects.

Third, this study focuses on patients with drug-resistant epilepsy undergoing intracranial EEG monitoring. These patients typically exhibit more frequent and severe seizures, and their medication regimens are more complex compared to patients undergoing non-invasive monitoring. However, our application of interest is with implantable devices, for which drug-resistant patients are the most common candidates. Future work may expand these analyses to include scalp EEG data and a wider range of epilepsy types to validate these findings in more diverse clinical settings.

Lastly, it is possible that ASM load modulates interictal spike rate, which consequently affects synchrony. While this study does not directly account for spike rates, prior research investigating the relationship between ASM load and spike rates has yielded conflicting results, suggesting substantial variability across patients. ^6,18,41,42^ Importantly, our findings demonstrate that synchrony consistently tracks with ASM load, independent of these unresolved questions about spike rate.

Our vision for clinical applications of our findings includes further studies to validate these measures in less invasive electrodes, small numbers of channels, and in patients who may not need epilepsy surgery. We envision an interactive system that allows for reciprocal communication between the patient and the wearable device interface. In such a system, synchrony may be only one of multiple measures—such as seizure counting, spike recording, sleep analysis—that may help patients with epilepsy actively manage and live with their disorders, rather than just being subject to unexpected events, such as seizures, drug reactions, toxicity and fluctuations for reasons that may not be apparent. We believe that such systems have tremendous potential to improve the lives of individuals with drug-resistant epilepsy, and perhaps other disorders, who cannot be “cured” by current therapy.

## Conclusion

This study provides compelling evidence that iEEG synchrony serves as a robust biomarker for ASM load. The significant negative correlation observed between ASM load and synchrony not only suggests that synchrony can track ASM levels but also aligns with the theoretical model of brain criticality, wherein ASMs maintain a subcritical state to prevent epileptic seizures. These findings carry important clinical implications, particularly for the development of implantable neurostimulation devices, which could leverage synchrony as a biomarker to enable real-time, patient-centred seizure management. Future research should focus on validating synchrony as a biomarker in outpatient settings with fewer electrodes, exploring its potential in broader epilepsy populations, and assessing its applicability across different EEG modalities. Additionally, the integration of synchrony monitoring into neurostimulation devices holds promise for shifting toward a more proactive, “patient-in-the-loop” seizure management paradigm, offering new opportunities for personalized epilepsy care.

## Supplementary material

Supplementary material is available at *Brain* online.

## Data availability

All iEEG recordings are available on https://www.ieeg.org/. Statistical analyses were performed using custom scripts implemented in Python and MATLAB with relevant packages and are available on GitHub (https://github.com/devincma/asm_synchrony_paper). Medication data is available upon request.

