## Supplementary Material for "Synchrony is a robust iEEG biomarker for antiseizure medication load in epileptic patients"

### Table S1 Patient demographics and epilepsy information.

| HUP number | Sex | Age at epilepsy onset | Age at implant | SOZ lateralization | SOZ localization | Engel outcome classification (1 year) | Had medication taper? | Had medication re-initiation? | Included in mixed model? |
|---|---|---|---|---|---|---|---|---|---|
| 137 | M | 36 | 53.2 | Left | Temporal | N/A | | | Y |
| 138 | M | 29 | 38.1 | Left | Temporal | IIA: Initially free of disabling seizures but has rare seizures now | | | Y |
| 139 | M | 0.03 | 19.8 | Left | Parietal | IB: Non disabling simple partial seizures only since surgery | | | Y |
| 140 | F | 26 | 47.4 | Left | Temporal | IB: Non disabling simple partial seizures only since surgery | Y | Y | Y |
| 141 | M | 14 | 30.1 | Right | Temporal | IIB: Rare disabling seizures since surgery | | | Y |
| 142 | M | 15 | 30.5 | Left | Parietal | IIIA: Worthwhile seizure reduction | Y | Y | Y |
| 143 | F | 14 | 29.2 | Right | Temporal | N/A | Y | | Y |
| 144 | M | 5 | 32.1 | Right | Frontal | ID: Generalized convulsions with antiepileptic drug withdrawal only | | | Y |
| 145 | M | 11 | 21.1 | Right | Temporal | IA: Completely seizure-free since surgery | Y | Y | Y |
| 146 | M | 4 | 19.5 | Bilateral independent Seizure Onsets | Frontal | IA: Completely seizure-free since surgery | | | Y |
| 147 | F | 12 | 44.7 | Left | Temporal | N/A | | | Y |
| 148 | M | 15 | 23.6 | Left | Temporal | IA: Completely seizure-free since surgery | | | Y |
| 149 | M | 43 | 49.7 | Right | Temporal | N/A | | | Y |
| 150 | M | 3 | 16.9 | Right | Temporal | IA: Completely seizure-free since surgery | | | Y |
| 151 | M | 6 | 33.6 | Right | Anterior cingulate | IA: Completely seizure-free since surgery | Y | | Y |
| 152 | F | 28 | 33.7 | Left | Multi-focal | N/A | Y | | Y |
| 153 | F | 43 | 56.3 | N/A | N/A | N/A | | | Y |
| 154 | F | 9 | 43.7 | Left | Frontal | N/A | | | Y |
| 155 | F | 4 | 38.6 | Left | Temporal | N/A | | | Y |
| 156 | F | 12 | 43.4 | Right | Temporal | N/A | | | Y |

| | | | | | | | | | |
|---|---|---|---|---|---|---|---|---|---|
| 157 | M | 16 | 25.6 | Left | Temporal | IIIA: Worthwhile seizure reduction | | | Y |
| 158 | M | 7 | 32.5 | Right | Frontal | IVA: Significant seizure reduction | | | Y |
| 159 | M | 17 | 22.7 | Left | Temporal | N/A | | | Y |
| 160 | F | 15 | 46.2 | Right | Temporal | IA: Completely seizure-free since surgery | Y | | Y |
| 161 | M | 40 | 59 | Right | Frontal | IVB: No appreciable change | | | Y |
| 162 | F | 22 | 35.4 | Left | Temporal | IB: Non disabling simple partial seizures only since surgery | | | Y |
| 163 | F | 36 | 41.3 | Left | Temporal | IIB: Rare disabling seizures since surgery | Y | Y | Y |
| 164 | F | 14 | 34.1 | Left | Temporal | IIA: Initially free of disabling seizures but has rare seizures now | Y | | Y |
| 165 | F | 11 | 20.6 | Right | Temporal | IIB: Rare disabling seizures since surgery | Y | | Y |
| 166 | M | 4 | 27.6 | Left | Temporal | IVA: Significant seizure reduction | Y | Y | Y |
| 167 | F | 2 | 40 | Generalized onset | Generalized | N/A | | | Y |
| 168 | M | 1 | 27.4 | Generalized onset | Generalized | N/A | | | Y |
| 169 | M | 20 | 23 | Left | Temporal | N/A | | | Y |
| 170 | M | 3 | 20.3 | Right | Insula | IVA: Significant seizure reduction | | | Y |
| 171 | M | 6 | 49.1 | Left | Frontal | IIIA: Worthwhile seizure reduction | Y | Y | Y |
| 172 | F | 2 | 28 | Left | Frontal | IB: Non disabling simple partial seizures only since surgery | | | Y |
| 173 | F | 12 | 23.8 | Right | Temporal | IA: Completely seizure-free since surgery | | Y | Y |
| 174 | F | 16 | 22.2 | Left | Non-localizable | N/A | | | Y |
| 175 | F | 6 | 20 | Generalized onset | Generalized | N/A | | | Y |
| 177 | F | 5 | 42.4 | Right | Temporal | IA: Completely seizure-free since surgery | Y | Y | Y |
| 178 | F | 28 | 38.2 | Right | Temporal | IIB: Rare disabling seizures since surgery | | | Y |
| 179 | F | 13 | 19.8 | Left | Frontal | ID: Generalized convulsions with antiepileptic drug withdrawal only | | | Y |

| | | | | | | | | | |
|---|---|---|---|---|---|---|---|---|---|
| 180 | F | 2 | 28.1 | Left | Frontal | IA: Completely seizure-free since surgery | | | Y |
| 181 | F | 16 | 31 | Left | Temporal | IA: Completely seizure-free since surgery | | | Y |
| 182 | F | 19 | 25.6 | Left | Temporal | N/A | Y | | Y |
| 184 | M | 18 | 22.5 | Left | Temporal | IB: Non disabling simple partial seizures only since surgery | | | Y |
| 185 | M | 9 | 39.4 | Left | Temporal | IA: Completely seizure-free since surgery | | | Y |
| 186 | F | 15 | 36.2 | Left | Temporal | N/A | | | Y |
| 187 | M | 18 | 24.4 | Right | Temporal | IB: Non disabling simple partial seizures only since surgery | | | Y |
| 188 | F | 10 | 24.5 | Left | Frontal | IVB: No appreciable change | | | Y |
| 189 | M | 28 | 29.4 | Left | Temporal | IVB: No appreciable change | | | Y |
| 190 | M | 9 | 24.8 | Left | Temporal | IA: Completely seizure-free since surgery | Y | Y | Y |
| 191 | F | 16 | 32.7 | Left | Temporal | IIA: Initially free of disabling seizures but has rare seizures now | | | Y |
| 192 | F | 23 | 46.7 | N/A | N/A | N/A | Y | | Y |
| 193 | M | 6 | 42.3 | N/A | N/A | N/A | | | Y |
| 194 | F | 7 | 48.2 | N/A | N/A | IB: Non disabling simple partial seizures only since surgery | | | |
| 195 | F | 25 | 27.8 | Left | Temporal | N/A | | | |
| 196 | F | 3 | 58.8 | Inconclusive | Non-localizable | N/A | | | Y |
| 197 | F | 1 | 40.3 | Left | Temporal | N/A | | | |
| 199 | F | 8 | 44.8 | Bilateral independent Seizure Onsets | Temporal | N/A | | | Y |
| 201 | F | 16 | 37 | Left | Temporal | IB: Non disabling simple partial seizures only since surgery | | | |
| 202 | F | 38 | 39.8 | Left | Temporal | N/A | | | Y |
| 204 | M | 28 | 29.9 | Left | Temporal | IIB: Rare disabling seizures since surgery | | | Y |
| 205 | F | 13 | 23.2 | Right | Temporal | N/A | | | Y |
| 206 | F | 6 | 47.4 | Right | Temporal | IID: Nocturnal seizures only | | | Y |
| 207 | M | 21 | 40 | Left | Temporal | IA: Completely seizure-free since surgery | | | Y |
| 208 | M | 16 | 28.1 | Left | Multi-focal | IVA: Significant seizure reduction | | | |

| | | | | | | | | | |
|---|---|---|---|---|---|---|---|---|---|
| 209 | F | 18 | 40.7 | Left | Temporal | N/A | | | |
| 210 | M | 12 | 29.4 | Left | Frontal | N/A | | | Y |
| 211 | M | 12 | 23.3 | Bilateral independent Seizure Onsets | Temporal | N/A | | | |
| 213 | M | 2 | 21.1 | Left | Frontal | N/A | | | |
| 214 | M | 32 | 40.6 | Bilateral independent Seizure Onsets | Temporal | N/A | | | |
| 215 | M | 30 | 36.6 | Left | Multi-focal | ID: Generalized convulsions with antiepileptic drug withdrawal only. | | | |
| 216 | F | 3 | 21.1 | Left | Multi-focal | IA: Completely seizure-free since surgery | | | |
| 217 | F | 16 | 39.2 | Right | Temporal | IA: Completely seizure-free since surgery | | | Y |
| 219 | M | 22 | 26.8 | Right | Temporal | IIC: More than rare disabling seizures after surgery, but rare seizures for at least 2 years | | | Y |
| 221 | M | 1 | 38.3 | Right | Temporal | IA: Completely seizure-free since surgery | | | Y |
| 223 | M | 37 | 46.6 | Left | Temporal | IB: Non disabling simple partial seizures only since surgery | | | Y |
| 224 | F | N/A | 42.8 | Inconclusive | Non-localizable | N/A | | | |
| 225 | M | 13 | 33 | Right | Temporal | IA: Completely seizure-free since surgery | | | Y |
